# Collective aggressiveness limits colony persistence in high but not low elevation sites in Amazonian social spiders

**DOI:** 10.1101/610436

**Authors:** James L. L. Lichtenstein, David N. Fisher, Brendan L. McEwen, Daniel T. Nondorf, Esteban Calvache, Clara Schmitz, Jana Elässer, Jonathan N. Pruitt

**Author notes:** Denotes corresponding author.

## Abstract

Identifying the traits that foster group survival in contrasting environments is important for understanding local adaptation in social systems. Here we evaluate the relationship between the aggressiveness of social spider colonies and their persistence along an elevation gradient using the Amazonian spider, *Anelosimus eximius*. We found that colonies of *A. eximius* exhibit repeatable differences in their collective aggressiveness, and that colony aggressiveness is linked with persistence in a site-specific manner. Less aggressive colonies are better able to persist at high-elevation sites, which lack colony-sustaining large-bodied prey, whereas colony aggression was not related to chance of persistence at low-elevation sites. This suggests resistance to resource limitation through docility promotes colony survival at high elevations. These data reveal that the collective phenotypes that relate to colony persistence vary by site, and thus, the path of social evolution in these environments is likely to be affected.

## Introduction

Although social evolution provides numerous benefits for group constituents (Krause & Ruxton, 2002), social groups can also vary considerably in their success (ants: Gordon, 2013, social spiders: Aviles, 1986, honey bees: Watanabe, 2008). For a variety of social organisms, many or most of the social groups ever founded will swiftly end in their collective demise (Tibbetts & Reeve, 2003, Hahn & Tschinkel, 1997, Aviles & Tufino, 1998). In some taxa, even social groups in apparent good health can fall victim to colony extinction events (Pruitt, 2012). Thus, any feature that enables groups to persist in their environment is likely to foster their success. Social organisms provide an interesting case study for evolutionary ecologists, because trait differences occur at both the individual level and between groups, in terms of their collective traits (Jandt et al., 2014, Bengston & Jandt, 2014, Wray & Seeley, 2011). Like individual traits, a growing body of evidence conveys that group traits are often associated with group success (Shaffer et al., 2016, Gordon, 2013, Wray et al., 2011), and that these links can vary between environments (Pruitt & Goodnight, 2014, Pruitt et al., 2018). Site-specific selection may therefore contribute to biodiversity by promoting intraspecific variation and local adaptation in group-level traits.

Social spiders are a useful model with which to explore the evolutionary ecology of group extinction events and collective behavior in general. This is because social spider groups emerge and disappear with high frequencies (reviewed in Aviles & Guevara, 2017). This, and because groups are inbred and composed of highly related individuals (Riechert & Roeloffs, 1993, Aviles, 1993, Henschel et al., 1995), means that group success is a major determinant of individuals’ inclusive fitness. Here we explore the degree to which group behavior is linked with group persistence using a highly social spider, the Amazonian spider *Anelosimus eximius* (Araneae, Theridiidae). This species occurs across a range of habitat types from Panama to Argentina at varying elevations. We use this variation in elevation to examine whether the relationship between group behavior and persistence varies along an elevation gradient. In particular, we hypothesise that collective aggressiveness should be favored at sites with low prey availability (Pruitt et al., 2018). For *A. eximius*, high-elevation sites are reasoned to be resource-limited because they harbor smaller average prey sizes (Yip et al., 2008, Powers & Aviles, 2007, Guevara & Aviles, 2007, Guevara & Aviles, 2015). By contrast, we predict that less aggressive colonies will be favored in high-resource and enemy-rich environments, like lowland rainforests (Purcell & Aviles, 2008). Thus, we predict that selection on collective aggressiveness will mimic the usual patterns observed in solitary spiders and other taxa, where low resources favor heightened aggression and responsiveness towards prey (Riechert, 1993, Magurran & Seghers, 1991, Dunbrack et al., 1996). If this is so, then it would hint that theory developed for behavioral evolution in solitary organisms can be redeployed to correctly predict patterns of selection occurring at the level of collective traits.

## Materials and Methods

### Focal species and sites

We measured collective foraging aggressiveness in colonies of *A. eximius* across the Ecuadorian Amazon in Oct.-Nov. 2017. *A. eximius* colonies build basket-shaped nests with large capture webs where they hunt collectively. We observed colonies at three sites on the e45 near Archidona (n=14; S 0° 46.214, W 77° 46.604), the e20 towards Coca (n=10; S 0° 43.421, W 77° 39.993), and near the Iyarina lodge (n=9; S 1° 4.027, W 77° 37.228). We further sampled two sites: roadsides, forest interiors, and waterways in the Yasuní National Park (n=16; S 0° 40.862, W 76° 23.152) and waterways near the Cuyabeno Wildlife Reserve (n=21; S 0° 1.921, W 76° 12.851).

### Collective aggressiveness

We measured colonies’ aggressiveness by placing dummy prey (1cm sections of dead leaf) 4cm from the rim of the nest basket, and vibrating it with a handheld vibratory device until spiders emerged and seized the dummy prey (Pruitt et al., 2017), between 1000-1600 hours. We recorded the latency of the first spider to contact the dummy. We subtracted the attack latency from 600 to obtain an aggression index where higher scores correspond to higher aggressiveness. We repeated these tests every day for four days on a subset of colonies at Archidona (n=11), Iyarina (n=4), and Yasuní (n=10), to assess the repeatability of colony aggressiveness. For the remaining colonies, aggressiveness was only measured once due to logistical constraints. Latency to attack prey is a common measure of foraging aggressiveness in solitary and social spiders (Riechert & Hedrick, 1993, Pruitt et al., 2013, Kralj-Fiser & Schneider, 2012, Kralj-Fiser et al., 2012), and it tightly linked with prey capture success and foraging performance in several species of group-living spiders (Kamath et al., 2018, Pinter-Wollman et al., 2017, Pruitt & Riechert, 2011).

### Habitat measurements and persistence

Immediately following aggressiveness assays, we also recorded habitat characteristics and marked colonies with aluminium tree tags. First, we recorded colony elevation and GPS coordinates (Garmin eTrex 30x). Then, the canopy cover over each colony was estimated with using the iPhone application Canopyapp (Davis et al., 2018). We assessed carnivorous ant presence by measuring latency of ant recruitment to 35g of tuna within 2m of the web (Hoffman & Avilés, 2017), placed on the forest floor beneath the colony. A subset of colonies was run through two such ant-baiting tests, and microhabitat differences in ant recruit speed were found to be consistent through time even within a specific site (r = 0.86, 95% CI: 0.57-0.96, p < 0.0001, n = 21). Faster ant recruitment times were taken as evidence that the microhabitat immediately around the focal colony had a greater risk of attack by predatory ants.

We estimated the volume of web baskets by measuring the size of the smallest possible orthotope that contained the basket, by first approximating the shape of each web (e.g., square base, circle base) and then taking the necessary measurements to compute the web volume. Web volume increases approximately linearly with group size in *A. eximius* (Yip et al., 2008, Powers & Aviles, 2007). To determine colony survival, we returned in Oct. 2018, eleven months later, and recorded whether the colony contained any remaining living individuals. This time interval corresponds to ∼2 generations of *A. eximius* (Vollrath, 1982). All aluminum tags were then removed.

### Statistical methods

We could not satisfactorily fit a generalised linear model simultaneously evaluating the influence of elevation, aggression and colony size on persistence. Moreover, neither colony aggression nor elevation could satisfactorily be transformed towards normality. Finally, aggressiveness was not repeatable within sites, r = 0 (95% CI: 0.0 - 0.157, p = 0.500), indicating that colonies’ behavior within each site are relatively independent. Therefore, we compared the elevation, aggressiveness, and web size of colonies that either persisted or not using Mann-Whitney U-tests. We assessed the correlation between elevation and aggressiveness, and aggressiveness and colony size using Spearman rank correlations. We took the log of basket volume as our index of colony size.

To determine whether the relationship between colony persistence and aggression depended on the elevation of the colony, we split the data into “high” elevations (above 740m, 25 colonies) and “low” elevations (below 450m, 43 colonies). This split demarcates a natural break in our sampling distribution. We then compared the aggressiveness of colonies that persisted or not in each dataset separately using Mann Whitney-U tests. To determine how canopy cover and the presence of predator ants varied with elevation, we performed Spearman rank correlations between elevation and each of canopy cover and the latency for ants to arrive at the tuna bait. There were 71 focal colonies in total. However, three colonies did not have elevations recorded. Four colonies had no web size measurements, owing to their residing in relatively inaccessible microhabitats (e.g., suspended over cliffs). Otherwise, sample sizes for each group in each comparison are given below. The repeatability of colonies’ aggressiveness was assessed by fitting linear a mixed model with “aggressiveness” as the response variable, “colony ID”, “site”, and “trial iteration”, using the rptR package (Stoffel et al., 2017). This allows us to estimate the intra-class correlation coefficient of colony ID, while accounting for variance explained by site and trial iteration. We estimated 95% confidence intervals on repeatability estimates by running the linear mixed model though 1000 bootstrap iterations. As mentioned above, we aimed to measure 25 colonies across three sites four times each, although three colonies only received three measurements, giving 97 measurements across 25 colonies in total to assess repeatability.

## Results

The influence of aggression on persistence depended on altitude. At high elevations, persisting colonies were less aggressive (mean = 505, n = 19) compared to colonies that vanished (mean = 592, n = 6; Fig. 1; Wilcox test, W = 2, p < 0.001). At low elevations, colonies that persisted were more aggressive (mean = 582, n = 27) than colonies that vanished (mean = 562, n = 16) but this difference was not significant (W = 272, p = 0.165). Although we could not satisfactorily fit a glmm to our data, the results of a glmm analysis qualitatively matched the results presented here (model predicting colony survival [*aggression x elevation*]: Est = −13.9 ± 6.30, z = −2.21, p = 0.027).

**Figure 1.**
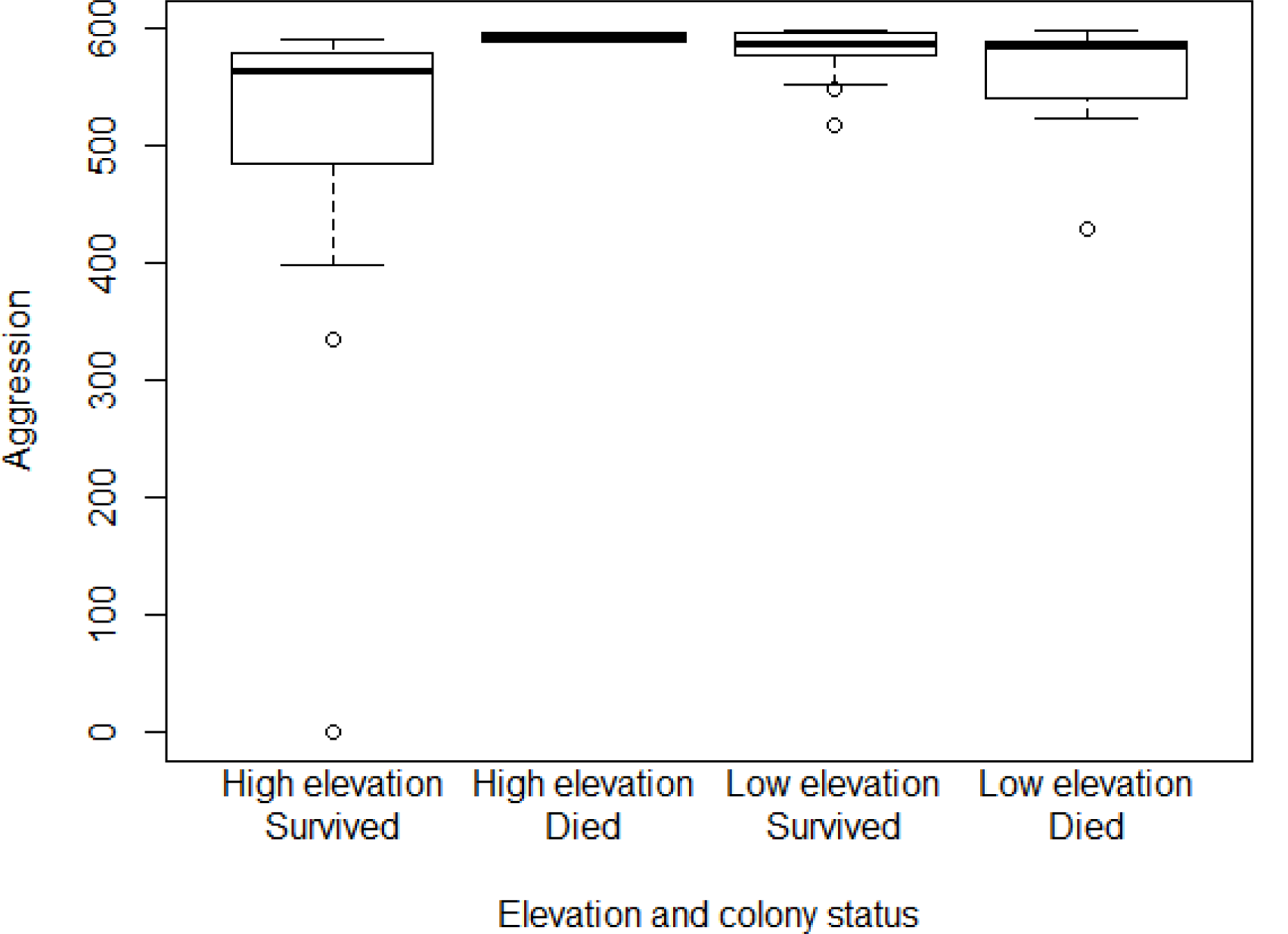
The aggressiveness of colonies that either survived or died, at low (< 450m) or high (>740m) elevation sites. Aggression was 600 minus the latency to attack (maximum 600 seconds) hence is unitless.

Elevation did not influence colony persistence. The mean elevation of colonies that persisted and vanished was 584m and 479m respectively (Fig. S1; n = 46 & 22 respectively, Wilcox test W = 570, p =0.404). Colony web size did not predict persistence; colonies that persisted were no larger than those than did not. Medians (means are highly biased by a few large value) of volume were 143,918 cm^3^ for colonies that persisted and 90,450cm^3^ for colonies that vanished, but the median logged values are 11.87 and 11.41 respectively (Fig. S1; n = 46 & 21 respectively, Wilcox test, W = 554, p = 0.344).

Colonies’ aggressiveness was not related to their web size (Fig. S2; n = 67, Spearman rank correlation, S = 47550, p = 0.691, rho = 0.051), but colonies were more aggressive at lower elevations (Fig. S2; n = 68, Spearman rank correlation, S = 65398, p = 0.041, rho = −0.248). Colony aggression was repeatable, r = 0.26 (95% CI: 0.012 - 0.474, p = 0.003).

Higher elevations were associated with reduced canopy cover (Spearman rank correlation, S = 66623, p = 0.006, rho = −0.329) and the slower recruitment of ants (Spearman rank correlation, S = 21568, p = 0.050, rho = 0.263).

## Discussion

Understanding the forces that enable some groups to persist and proliferate when others crash or disband is helpful for predicting how social evolution proceeds in contrasting environments. For many social animals, this can be thought of as a kind of group-level viability selection. Colonies of the Amazonian social spider *A. eximius* exhibit clinal variation in selection on aggressiveness. At odds with our *a priori* predictions, less aggressive colonies outperform their aggressive rivals at resource-poor high elevations. The opposite trend emerges at low elevations, although it was not statistically significant. Given this pattern of selection, one might predict that high elevation *A. eximius* should be less aggressive overall, either because of local adaptation or via on-going viability selection against aggressive colonies. Consistent with this prediction, we observed that colonies of *A. eximius* at higher elevation do indeed exhibit lower aggressiveness than their low-elevation counterparts. In aggregate, this conveys that site-specific selection on colony aggressiveness could play a role in generating geographic variation in colony behavior, akin to patterns observed in solitary species (Drummond & Burghardt, 1983, Magurran & Seghers, 1991, Riechert, 1993, Walsh et al., 2016).

The mechanisms underlying the success of non-aggressive colonies at high elevation remain elusive. We predicted that low-resource conditions would favor colonies with swifter foraging responses because, in trap-building predators, foraging is a time-sensitive opportunity. Thus, colonies at high elevations should maximize on the limited foraging opportunities that are available to them (Powers & Aviles, 2007, Guevara & Aviles, 2007). This is often the case for individual-level aggressiveness (Riechert, 1993, Magurran & Seghers, 1991, Dunbrack et al., 1996). However, it is perhaps equally plausible that low-resource conditions could favor reduced aggressiveness. If more aggressive colonies engage in more infighting, exhibit higher metabolic rates, or are otherwise more susceptible to starvation, then selection may favor less aggressive colonies under low resource conditions because it enables them to persist through times of resource scarcity. This mode of competition is often referred to as *Tilman’s R* Rule* (Tilman, 1982). Consistent with this hypothesis, there is evidence that both aggressive social *Anelosimus* (Lichtenstein & Pruitt, 2015) and *Stegodyphus* (Lichtenstein et al., 2017) are more susceptible to starvation, and that non-aggressive *Stegodyphus* colonies can outperform their rivals when resources fall below a critical level (Pruitt et al., in press). Alternatively, smaller average prey sizes at high elevation sites might merely not require the same levels of aggressiveness to subdue than the larger prey of low elevation sites. More detailed work within sites is needed to tease apart the mechanisms responsible for this among-site result.

We found that ants recruited more quickly to tuna baits at lower elevations. This suggests that the threat of predation from ants, or perhaps the degree of indirect resource competition from ants, will be higher at lower elevations. Either of these could select for higher aggressiveness (or, at least, against docility) in social spiders, which are more frequently attacked by ants a low-elevation sites (Purcell & Aviles, 2008, Hoffman & Avilés, 2017), and this may help to explain the patterns of selection that we observed. We also observed reduced canopy cover at higher elevations. While this seems unlikely to directly influence spider colony survival, it may influence the availability of prey (i.e. decreased cover may decrease the number of flying invertebrates) or increase web damage costs, and thus, have consequences for the benefits of colony aggression.

At odds with previous work, group size was not a significant predictor of colony persistence in our field data on *A. eximius*. The formation of larger coalitions is frequently associated with reduced group failure rate in social arthropods, and this fact is thought to underlie the formation of social life history trajectories like foundress coalitions in wasps and ants (Fewell & Page, 1999, Seppa et al., 2002, Tibbetts & Reeve, 2003, Miller et al., 2018). Group size dependent survival has also been documented in a number of social (Bilde et al., 2007, Aviles & Tufino, 1998) and transitionally social species of spiders (Lichtenstein et al., 2018). We reason that this discrepancy between findings is because colonies of the smallest size classes (one to a few dozen spiders) are largely missing from our data set, and the persistence benefits of increasing group size are most pronounced at the smallest colony sizes (Lichtenstein et al., 2018, Aviles & Tufino, 1998).

In summary we detected a site-specific relationship between colony aggressiveness and persistence in a social spider. Furthermore, we found a cline in aggression with elevation that suggests that the selective benefits to reduced aggression at higher elevations are strong enough to promote appropriate fit between colony traits and the habitats in which they reside.

## Ethics

The studies herein were conducted on invertebrates and were therefore not subject to ethics approval. Field studies were conducted under research permit N°23-17 IC-FAU-DNB/MA.

## Data accessibility

The data for this manuscript can be found on Dryad: https://datadryad.org/review?doi=doi:10.5061/dryad.hr90jf2

## Authors Contributions

JLLL and BLM assisted with all aspects of the study pipeline. DTN, EC, CS and JE assisted with data collection. DNF and JNP helped to analyze the data and write the paper.

## Acknowledgements

We are indebted to the Ecuadorian ministry of the environment for granting our research permit (N°23-17 IC-FAU-DNB/MA) and Dr. Clifford Kiel for his sponsorship. We would like to thank the Yasuní Scientific Station of the Pontifical Catholic University of Ecuador and Tod Swanson from the Andes and Amazon Field School for logistical assistance in the field.

## Figures & Supplementary Figures

**Figure S1.**
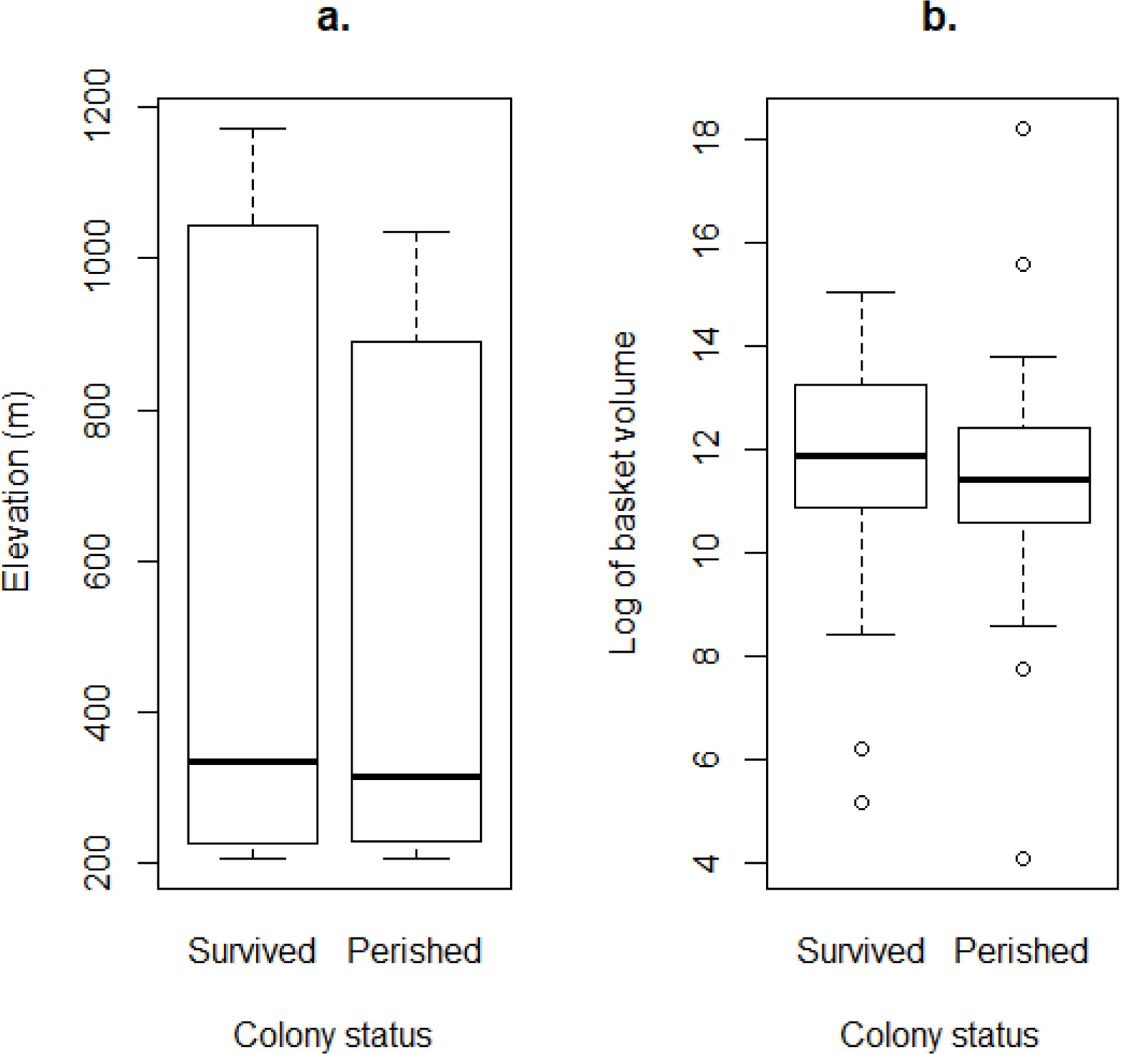
The difference in elevation (metres, a.) and colony size (the log the basket volume, b.) of colonies that either survived or perished. Neither elevation nor colony size differed between colonies that survived or perished.

**Figure S2.**
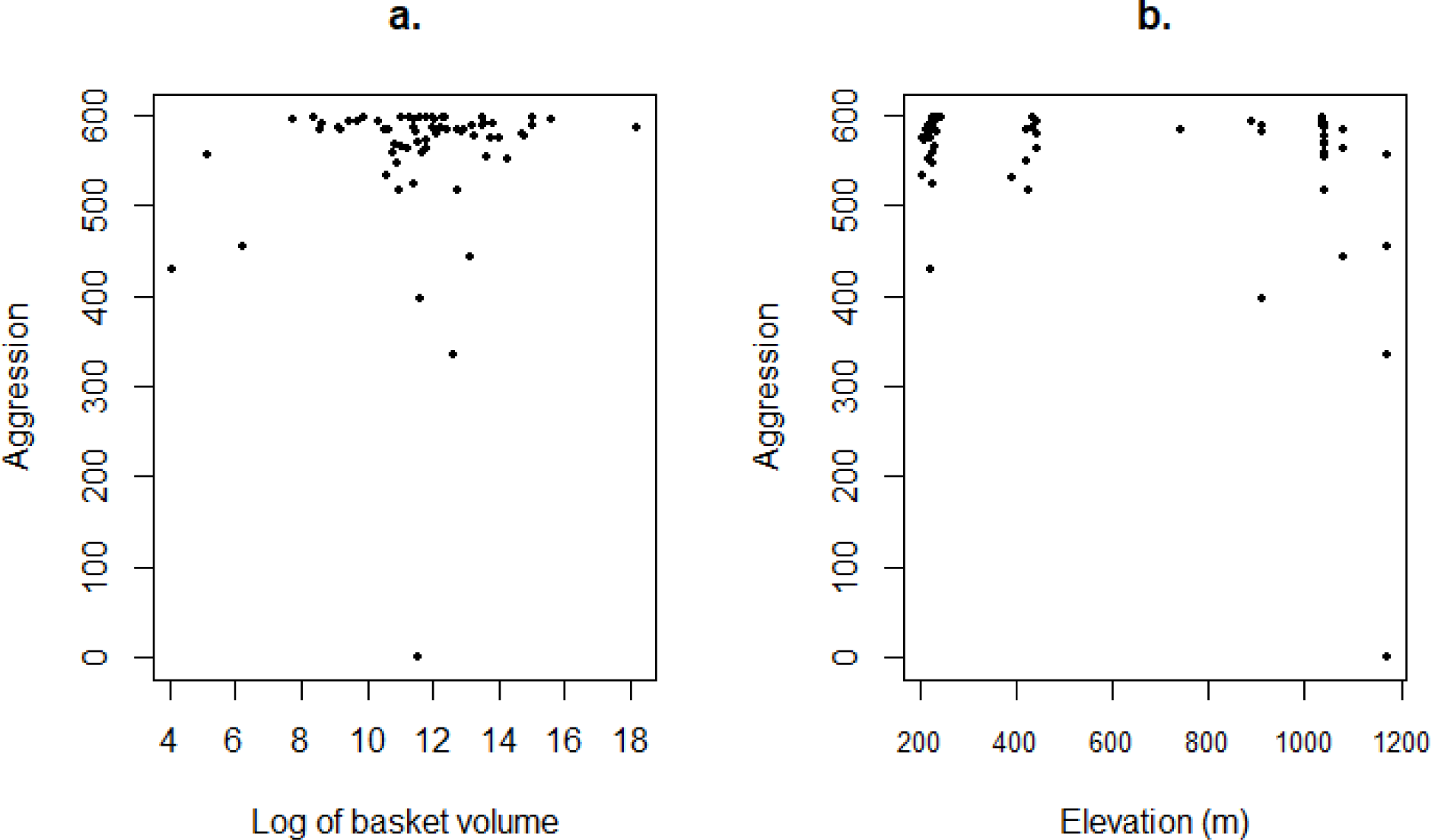
The relationship between colony aggression and colony size (log of basket volume, a.), and elevation (metres, b.). Aggression was not related to colony size, while it is weakly negatively correlated with elevation.

